# Seeing more than schemas: the vmPFC represents imagery-rich mental scenarios

**DOI:** 10.1101/2025.08.01.667951

**Authors:** Sorit Achmed Ali, Pitshaporn Leelaarporn, Rüdiger Stirnberg, Maren Bilzer, Nadja Abdel Kafi, Julia Taube, Yilmaz Sagik, Cornelia McCormick

**Author notes:** **Corresponding author:** Cornelia McCormick Department for Old Age Psychiatry and Cognitive Disorders Universitätsklinikum Bonn Venusberg-Campus 1 53127 Bonn.

## Abstract

Mental imagery varies dramatically across individuals, from vivid scene construction to the complete absence of visual experience, as seen in aphantasia. While the ventromedial prefrontal cortex (vmPFC) is traditionally associated with abstract, schematic representations, emerging theories suggest it may also contribute to constructing vivid, visual mental content. To test this, we developed a novel 7T fMRI experiment varying imagery demands across conditions: Prior to scanning, participants memorized richly detailed scenarios, more constrained, stationary objects, and finally semantic definitions for each of three abstract German words (e.g., hope). During fMRI and eye-tracking, the same word was presented across trials, but participants vividly re-engaged with one of three distinct representations (i.e., scenarios, objects, and definitions), allowing for comparison across richly different cognitive modes triggered by identical visual input. Univariate analyses confirmed previous findings; highlighting the roles of the vmPFC, hippocampus, parahippocampal cortex, and visual-perceptual cortex in imagery-rich scenario construction. We further performed multivoxel pattern analysis (MVPA) to examine distributed neural representations, which can reveal the informational content of mental imagery beyond activation magnitude. Critically, the vmPFC was the only brain region where MVPA classifier accuracy was higher for scenario construction than for object and abstract conditions, directly supporting our hypothesis that the vmPFC encodes imagery-rich details rather than solely abstract, schematic information. Eye movement variability also distinguished between conditions. These findings advance our understanding of vmPFC function, emphasising its role in representing vivid mental content.

**Highlights:**

- · vmPFC distinctly represents imagery-rich scenarios in MVPA analysis
- Univariate fMRI reveals increased HC and PHC activity during scenario construction
- Eye movements vary by imagery type, distinguishing scenarios from objects and definitions

## 1. Introduction

Some people can vividly imagine a beach at sunset; others, like those with aphantasia, report seeing vaguely to nothing at all. Those with absent imagery not only report less vivid and detailed mental imagery in their subjective descriptions but also exhibit selective impairments in objective imagery tasks, such as binocular rivalry (Bainbridge et al., 2021, Monzel. Leelaarporn et al. 2024). This variability in mental imagery raises fundamental questions about how the brain constructs internal representations; particularly the role of regions like the ventromedial prefrontal cortex (vmPFC). Traditionally linked to abstract schemas, subjective value and reward, and self-referential processes(Delgado et al., 2016; Ghosh & Gilboa, 2014; Gilboa & Marlatte, 2017), the vmPFC is now increasingly associated with constructing vivid, perceptually rich mental scenarios(Ciaramelli et al., 2019; Kindler et al., 2025, preprint; McCormick, Ciaramelli, et al., 2018; Taube et al., 2025, preprint). This shift prompts a key question: does the vmPFC merely coordinate conceptual frameworks, or does it also neurally represent vivid imagery-rich thought?

Mental imagery underpins a range of cognitive functions including autobiographical memory, spatial navigation, future thinking, and mind-wandering (Hassabis et al., 2007; Maguire & Mullally, 2013; McCormick, Rosenthal, et al., 2018). In aphantasia, we have recently shown that autobiographical memory recall is less vivid and neurally supported differently (Monzel, Leelaarporn et al., 2024). Mental imagery is supported by a distributed network including the vmPFC, hippocampus (HC), parahippocampal cortex (PHC), superior temporal gyrus (STG), and visuoperceptual cortices within the occipital cortex.

Traditionally, the vmPFC has been associated with schema processing, emotion regulation, moral and economic decision-making, as well as social cognition (A Bechara, 2000; Delgado et al., 2016; Ghosh & Gilboa, 2014; Koenigs et al., 2008; Lockwood et al., 2024). However, emerging evidence suggests that it also plays a key role in initiating and coordinating vivid, scene-based mental imagery (Ciaramelli et al., 2019; Lieberman et al., 2019; McCormick, Ciaramelli, et al., 2018). A hierarchical model proposes that while the hippocampus constructs individual scenes, the vmPFC orchestrates broader temporally extended mental scenarios by guiding transitions between scenes through iterative interactions with the HC and neocortical areas (McCormick, Ciaramelli, et al., 2018; Taube et al., 2025, preprint). MEG studies support this framework, showing vmPFC activation precedes hippocampal engagement during scene construction and autobiographical memory retrieval (McCormick et al., 2020; Monk et al., 2021, Monk et al. 2020).

In fact, Monk (2020) proposed a top-down mechanism in which the vmPFC activates scene schemas and drives conceptual elaboration in the STG, a region acting as an amodal semantic hub (Chadwick et al., 2016). The STG, part of the anterior temporal lobe (ATL), is essential for integrating and storing conceptual knowledge (Pobric et al., 2007). Consistent with this view, (McCormick & Maguire, 2021) proposed that vmPFC connectivity shifts flexibly depending on task demands; strengthening with lateral temporal cortex during semantic processing and with HC and visual areas during spatially coherent scene construction. Despite these advances, prior research often confounds visual vividness with conceptual complexity. As a result, it remains unclear whether the vmPFC merely supports abstract, schematic reasoning, or if it is also preferentially involved in perceptually rich scene/scenario imagery. This question is particularly relevant to conditions like aphantasia, where people lack voluntary visual imagery but retain semantic knowledge.

In addition, the hippocampus plays a central role in binding diverse elements, such as objects, spatial layouts, and sensory features into coherent naturalistic scenes (Hassabis & Maguire, 2009; McCormick et al., 2021). Scenes are typically defined as egocentric, 3D, spatially coherent representations with multiple elements (Dalton et al., 2018; Lee et al., 2005). These internal models support episodic memory, imagination, and navigation (Hassabis et al., 2007; Maguire et al., 2016; Maguire & Mullally, 2013). HC activity is heightened during scene construction compared to object imagery or abstract reasoning (Clark et al., 2018), though emerging findings suggest that hippocampal subcircuits may be flexibly recruited based on task type (Dalton et al., 2018). As an adjacent brain region, the PHC supports contextual and spatial layout information (Aminoff et al., 2013; Epstein, 2008; Lee et al., 2005; Staresina & Davachi, 2010).

Visual cortex is involved in both object and scene imagery, but with different patterns. Early occipital areas (V1–V3) support reactivation of perceptual detail during vivid scene construction (Cui et al., 2007; Naselaris et al., 2015), while object imagery involves the lateral occipital complex (LOC), which represents form and shape (Emberson et al., 2017; Kim et al., 2009).

To investigate this question, we designed a novel 7T fMRI experiment in which participants, prior to scanning, memorized a scenario, a symbolic object, and a conceptual definition for each of three abstract German words (e.g., hope). During the eye-tracking and scanning experiment, the same word was displayed repeatedly across trials, prompting participants to vividly re-engage with one of the three learned representations, scenarios, objects, or definitions. This manipulation enabled us to systematically vary imagery demands, from richly detailed scenarios to more constrained, stationary objects, and finally to semantic definitions largely devoid of perceptual content. To examine the neural representations associated with each task, we applied multivoxel pattern analysis (MVPA), a technique that goes beyond univariate activation levels by capturing distributed patterns of activity, offering insights into the content and specificity of mental representations—including the degree of visual detail (Norman et al., 2006). Additionally, eye-tracking may capture oculomotor patterns that reflect internal scene construction process. Therefore, we conducted an eye-tracking experiment to assess condition-specific differences in visual attention strategies during imagery.

We hypothesized that the vmPFC would show represent vivid scenarios to a greater extent as abstract definitions or single objects. The hippocampus and PHC were expected to support scenario and object construction, while the STG was predicted to act as a semantic hub for the abstract definitions. Early visual areas were expected to contribute to perceptual richness in scenario and object construction. In eye-tracking, we anticipated higher fixation and saccade counts for scenarios, and longer fixation durations during object imagery.

## 2. Methods and Materials

### 2.1 Participants

26 young healthy, right-handed participants were included. Six participants were excluded due to technical issues during fMRI scanning. Thus, the final analysis was conducted with the data of 20 participants (11 females, 9 males) in the age range of 20-27 (mean age: 22,55 ± 1,60 years). Participants had no history of neurological or psychiatric disorders. At the time of the experiment, they did not experience depressive symptoms (mean 14.5 ± 8.9, Beck’s Depression Inventory (BDI-V) (Schmitt et al., 2006). Of note, only individuals with a BDI-V score of 35 or below were eligible but none were excluded based on this criterion. Since we aimed to assess vivid visual imagery, Aphantasia was an exclusion criterion. This was assessed using the Vividness of Visual Imagery Questionnaire (VVIQ) (Marks, 1973), where participants scoring above 32 were considered eligible. Our participants had a mean of 64.1 ± 12.1 VVIQ score and no one was excluded based on this criterion. All participants had normal or corrected-to-normal vision. Informed consents, both oral and written, were obtained from all participants, and the study was conducted in accordance with guidelines set forth by the local ethics board.

### 2.2 General experimental procedure

The study consisted of two study visits for each participant with an interval of 1-2 weeks in between. During the first visit, consent was obtained and all inclusion criteria were carefully checked. Afterwards, participants were familiarized with the experimental task and the training schedule was explained to them. Since MVPA examines overlapping neural activation for repeated presentation of the stimuli, much care was given to train participants before the fMRI scanner session. During the second visit, the experimenter ensured that the training was successfully completed. Afterwards, participants first completed the eye-tracking and then the fMRI experiment.

### 2.3 Imagery task during eye-tracking and fMRI

Our main goal was to investigate whether the vmPFC would represent vivid mental imagery over and above semantic scaffolding. Thus, we developed a novel experimental task in which we varied imagery demands across conditions. We asked participants to vividly imagine three abstract German words: *Hope* (*Hoffnung*), *Secret* (Geheimnis), and *Order* (*Ordnung*) either as a scenario, an object, or as an abstract definition.

For the scenario condition, participants were asked to vividly imagine a sequence of scenes lasting 7 seconds (i.e., trial duration). These scenarios were designed to be detailed, vivid, and contextually tied to the abstract word. For the word “hope”, the provided scenario was: *“I am at an auction and outbidding everyone there to get the painting that is up for grabs. I am full of hope and confidence that I will get it.”* The participants were asked to learn these scenarios by heart and imagine it as vividly as possible in front of their mind’s eye for the length of the trail duration.

For the object condition, participants were instructed to visualize a static object against a blank background with as much detail and clarity as possible. Dynamic or moving elements were explicitly discouraged. For the word *“hope”*, participants were instructed to imagine a stereotypical “white dove” -like line drawing without a background. The dove was chosen because of its strong symbolic association with hope. The participants were asked to learn the objects by heart and imagine this stationary object in front of their mind’s eye for the length of the trail duration.

For the abstract definition condition, we constructed concise definitions consisting of two to three sentences for each stimulus designed to take approximately 7 seconds to complete. The definitions were carefully formulated to maintain purely abstract conceptual content, systematically avoiding concrete or visually-evocative language. For instance, the definition for the word *“hope”* was: *“Hope means confidence and optimism. There is an expectation that good, positive things will happen in the future. The opposite is hopelessness and despair.”* Participants were asked to learn those definitions by heart and to recite these mentally without verbalizing.

Each of the three conditions was crossed with all three target words, yielding nine distinct stimuli, with standardized task instructions provided for participants to memorize.

### 2.4 Eye-tracking data acquisition

Before the eye-tracking experiment, participants were asked whether they adhered to the training schedule and were able to produce the scenarios, objects, and definitions accurately.

Eye movements were recorded using a video-based eye-tracker (EyeLink 1000, SR Research) at a sampling rate of 2000 Hz. The system tracked participants’ eyes while their head position was stabilized using a chin rest positioned 64 cm from the 47 cm wide screen display and 58 cm from the desktop-mounted camera. Stimuli were presented on slides displayed within the eye-tracker setup, using black Arial font on a dark gray background to enhance contrast and to avoid afterimages in the blank display. Participants were initially familiarized with the procedure through a brief instructional presentation, featuring slides identical to those used in the experiment.

Prior to the experiment, ambient lighting was turned off to minimize external visual distractions. Before the experiment started, a 9-point calibration and validation procedure was performed to ensure accurate tracking (average error < 0.5° of visual angle). Following calibration, the slide show commenced with an introductory instruction slide.

Each trial began with a jitter, which set the baseline. It was shown for 1 second. This was followed by a task slide, displaying a word and its associated condition for another second (e.g., “Hope”, “Scenario”). Subsequently, participants viewed a blank screen for 7 seconds, during which they mentally engaged in the imagery task. No specific instructions were given regarding their eye-movements. The task was succeeded by two questions. One of the questions targeted the success of the imagination. Participants should rate on a 4-point Likert scale, ranging from very good to very bad (i.e., 1=very good). The second question concerned the vividness of the imagination. Participants were again asked to rate on a 4-point Likert scale, ranging from very vivid to very faint (i.e., 1=very vivid). Important to note for the condition DF, the answer option for most successful trial for the question “vividness” was 4. Both questions were self-paced (up to a maximum of 3 sec). The Eye-tracking experiment was built using SR Research Experiment Builder v. 1.1 and consisted of 27 trials in total. This way each of the 9 tasks was repeated three times which lasted approximately 15 minutes.

### 2.5 MRI experiment

#### 2.5.1 MRI acquisition

Structural and functional MRI data were acquired using a MAGNETOM 7 T Plus ultra-high field scanner (Siemens Healthineers, Erlangen, Germany). We acquired a T1w-image, functional MRI data and a T2w-image (which is not part of the current manuscript).

#### T1-weigthed structural MRI

A whole-brain T1-weighted multi-echo MP-RAGE scan was acquired with 0.6 mm isotropic resolution using a custom sequence optimized for efficiency and minimal geometric distortions (Brenner et al. 2014). The scan parameters included an inversion time (TI) of 1.1 s, repetition time (TR) of 2.5 s, echo times (TEs) of 1.84, 3.55, 5.26, and 6.97 ms, flip angle (FA) of 7°, and a total acquisition time (TA) of 8:05 minutes. The readout pixel bandwidth was 970 Hz, with a matrix size of 428 × 364 × 256, elliptical sampling, and sagittal slice orientation. The scan incorporated CAIPIRINHA parallel imaging with a 1 × 2z1 undersampling scheme and on-line 2D GRAPPA reconstruction, along with a turbo factor of 218. The four echo-time images were combined into a single high-SNR image using a root-mean-square method.

#### Functional MRI

A specialized multishot 3D echo planar imaging (EPI) sequence was used to achieve whole-brain imaging with high spatial resolution and sufficient signal-to-noise ratio (SNR) at 7 T (see (Leelaarporn et al., 2024; Stirnberg et al., 2017; Stirnberg & Stöcker, 2021). The imaging parameters included a TE of 21.6 ms, TRvol of 3.4 s, flip angle of 15°, matrix dimensions of 220 × 220 × 140, and a readout pixel bandwidth of 1136 Hz. Partial Fourier sampling (6/8) was applied along the primary phase-encoding direction, with slices oriented obliquely to the anterior-posterior commissure. This configuration allowed for a nominal voxel resolution of 0.9 mm isotropic. Several advanced features were incorporated to optimize the sequence: Skipped-CAIPI (3.1 × 7z2), an SNR-optimized 7-fold CAIPIRINHA undersampling method combined with interleaved 3-shot segmentation, was used and reconstructed online with 2D GRAPPA. For phase correction, a single externally acquired scan per volume replaced the standard integrated phase correction per shot, reducing scan time. Echo train lengths were varied to exclude only the latest echoes outside a semi-elliptical k-space mask, ensuring high-resolution imaging. Additionally, a slab-selective binomial-121 water excitation approach was employed instead of conventional fat saturation, further improving efficiency.

A short 2 min. practice run was completed prior to the main functional imaging. Each of the five functional sessions lasted approximately 6 minutes. To minimize artifacts from non-steady-state signals, the first five volumes (corresponding to the 17-second pre-trial waiting period) were excluded from analysis. Additionally, the first session included a two-echo gradient-echo field-mapping scan (3 mm isotropic) completed in 35 seconds. In each functional session, a maximum of 105 volumes was acquired.

#### 2.5.2 fMRI procedure

Prior to functional MRI data acquisition, room lights were dimmed, and participants underwent a short practice to get accustomed to the response boxes. The experiment consisted of five functional sessions. Each session comprised 27 trials, resulting in a total of 135 trials across five fMRI sessions, with 15 repetitions for each of the nine tasks. Each trial began with a jittered baseline slide, displayed for a variable duration of 1–4 seconds. This was followed by a cue slide, which presented the word and condition for the task (e.g. “Hope”, “Definition”). The cue slide remained on the screen for 7 seconds, during which participants executed their learned mental tasks. After completing the task, participants were prompted to rate the success of the trial within 5 seconds on 4-point Likert-scale from 1-4, 1 (very good), 2 (good), 3 (bad), 4 (very bad). Directly following the scan, participants were given a questionnaire in which they were asked to assess how their concentration levels changed over the course of the five sessions.

### 2.6 After-scan reproduction task

After completing the eye-tracking and fMRI experiments, participants were instructed to reproduce the definitions they visualized during the experiment and to provide detailed drawings of the objects and scenarios they imagined. This approach aimed to evaluate participants’ compliance with the practice regimen at home and their adherence to task instructions during the experiment.

To quantify responses, a quality judgement score on a 5-point scale was developed, with 5 indicating the highest level of compliance or accuracy. The scores were based on, for the DF condition, the number of correct words, for the OB and SCN condition, points were given for 4 questions and awarded 1 point per question: Can one depict what is drawn? Are details included in the drawing? Is the drawing consistent with the rest of the participant’s drawings or is this one especially good or bad, thus deviating from the rest? Can one see the participant’s effort? To minimize subjective bias, two independent assessors rated the responses. Inter-rater reliability was evaluated using Spearman’s correlation to determine the degree of alignment between the two sets of ratings.

### 2.7 Data analysis

#### 2.7.1 Behavioural analysis

Behavioural data, such as success, vividness ratings, and eye-tracking parameters were analyzed using separate repeated-measures (rm) ANOVAs with experimental condition (SCN, OB, DF) as within-subjects factor. For data collected during eye-tracking, we extracted standard parameters during the imagination period using the Eyelink Data Viewer (SR Research). The parameters were 1. Fixation Count, 2. Average Fixation Duration, 3. Saccade Count, and 4. Average Saccade Amplitude for each participant for each trial. If the one-way-RM-ANOVAs yielded significant effects between conditions (i.e., p< 0.05) we conducted post hoc comparisons using Bonferroni’s multiple comparison tests, considering p-values < 0.05 as statistically significant.

#### 2.7.2 MRI data pre-processing and univariate analyses

For the preprocessing, we used SPM12 on MATLAB. In the first step the anatomical and the functional scans obtained from each subject were reoriented to be in line with the anterior-posterior commissure axis. The field maps, including the phase and magnitude images, were used to calculate the voxel displacement maps (VDM) to geometrically correct the distorted EPI images. The VDMs were then applied to the functional scans during realignment and unwarping. The structural T1w scans were co-registered with the functional scans and normalized to the Montreal Neurological Institute (MNI) space. The temporal high-pass filter was set to 128 s. Further, normalized functional data were smoothed using a Gaussian smoothing kernel of 6 mm full width half-maximum (FWHM).

To assess activity differences between our conditions, we performed a set of univariate contrasts. Three first level contrasts including motion regressors were specified: 1. SCN versus DF, 2. OB versus DF, and 3. SCN versus OB. These were then taken to the second level and thresholded at p<0.001 uncorrected and minimum cluster size of 10 contiguous voxels.

#### 2.7.3 Region-of-interest anatomical masks

We segmented 5 different ROI masks using the dimension corrected T1 structural image: bilateral vmPFC, bilateral hippocampus, bilateral occipital lobe, bilateral PHC and bilateral STG (see Fig. 1). To obtain the masks we used Fast-Surfer for automatic segmentation. Subsequently we applied manual correction using ITK-Snap for all ROIs. To apply bihemisperic analysis we collapsed the left and right mask for each ROI. For the vmPFC we orientated towards the region anterior to and below the genu of the corpus callosum (Delgado et al., 2016; McCormick, Ciaramelli, et al., 2018). We included areas of the orbitofrontal cortex (BA 11,12,13,14), of the frontal pole (BA 10), of subgenual cingulate cortex (BA 25) and of the anterior cingulate gyrus (BA 32) (Delgado et al., 2016; McCormick, Ciaramelli, et al., 2018). One participant had to be excluded from the vmPFC analyses because of signal drop out. The hippocampus mask encompasses the Uncus, the Dentate Gyrus, the Cornu Ammonis (1-4), the Subiculum and the Pre/Parasubiculum. The occipital mask covers the cuneus, the lingual gyrus, the pericalcarine area and parts of the fusiform gyrus towards the temporal border. The PHC lies adjacent to the hippocampus on the ventromedial edge of the temporal lobe. The STG constitutes the most superior gyrus of the temporal lobe.

**Figure 1.**
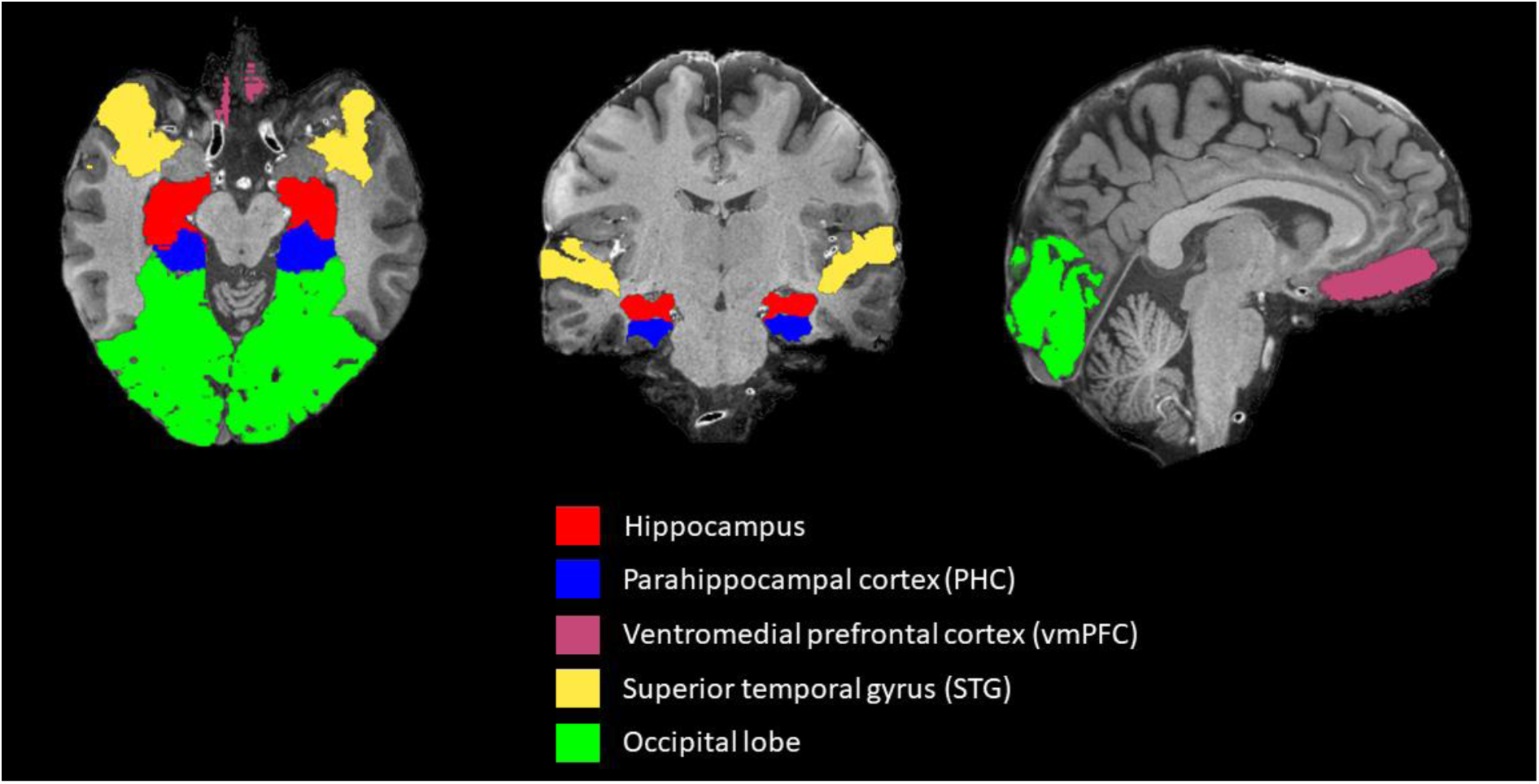
Example to anatomical region-of-interest masks overlaid on a single 7 Tesla T1w image in native space. All ROIs were extracted from individual segmentations using FastSurfer with additional manual correction.

Mean mask volumes (mean ± SD, in mm³) across participants were as follows: Hippocampus: 8625.5 ± 687.1 (one participant was excluded based on statistical outlier detection), Parahippocampal cortex (PHC): 6281 ± 720.6, Ventromedial prefrontal cortex (vmPFC): 12679361.3 ± 5540467.3, STG: 45382105.3 ± 5464184.9, Occipital Cortex: 123512500 ± 12409344.6. Volumes were calculated after binarization of masks.

#### 2.7.4 Multivoxel pattern analysis

To preserve spatial accuracy, native space fMRI data (no normalization, no smoothing) were used for MVPA analysis (see similar procedures in (Bonnici et al., 2012). Since three different cue words were used in three conditions, we calculated nine one sample t-test contrasts SCN1 vs Baseline (BL), SCN2 vs BL, SCN3 vs BL, DF1 vs BL, DF2 vs BL, DF3 vs BL, OB1 vs BL, OB2 vs BL, and OB3 vs BL. This procedure ensured that neural representations would be compared only to each other.

MVPA was then conducted using The Decoding Toolbox for MATLAB within SPM(Hebart et al., 2015). For classification, a support vector machine (SVM) was employed. The input data consisted of beta images derived from first level GLM analysis specified above. The classification procedure followed a leave-one-run-out cross-validation scheme, ensuring robust estimation of decoding accuracy across experimental runs. Both whole-brain and region-of-interest (ROI) analyses were performed. For all ROIs, we conducted bihemispheric analyses. For the ROI vmPFC, one participant was excluded due to incongruences between the ROI mask and the fMRI data.

The analysis output included 9×9 confusion matrices for each ROI, providing a detailed representation of classification accuracy across the nine conditions. We then averaged the accuracies of SCN1, SCN2, and SCN3 (and accordingly for OB and DF) into an averaged SCN classification accuracy. These classifier performance averages for all three conditions were summed and compared against chance level using one-tailed *t*-tests. Since we had three conditions, chance level was set to 33%. Following up, a one way-RM-ANOVA was conducted within the predefined ROIs to assess differences in classifier accuracy levels between the three experimental conditions. Subsequently, Tukey’s post hoc tests were conducted for multiple comparisons, with a significance threshold set at p <.05.

## 3. Results

### 3.1 Eye-tracking results

During eye-tracking, mean success ratings were M=1.47 (SD = 0.33) for SCN, M=1.29 (SD = 0.28) for OB, and M=1.50 (SD = 0.32) for DF. A significant main effect of Condition was observed, F(2, 18) = 13.194, p <.001. Post hoc analyses revealed that participants rated their imagination success significantly higher for SCN trials compared to OB trials (p <.001).

In addition, vividness ratings were highest for SCN (M = 1.66, SD = 0.41), followed by OB (M = 1.61, SD = 0.40), and lowest for DF (M = 3.34, SD = 0.83). As expected, vividness ratings differed significantly across conditions, F(2, 18) = 47.13, p <.001. Bonferroni-corrected post hoc tests showed that DF trials were rated significantly less vivid than both SCN and OB trials (p <.001).

Eye-tracking parameters revealed significant differences across conditions for fixation count F(2, 18) = 5.76, p =.012, average fixation duration F(2, 18) = 8.81, p =.002 and saccade count F (2,18) = 6.12, p =.009. No significant difference was observed for average saccade amplitude F(2, 18) = 2.63, p =.1.

Specifically, we found that fixation counts were highest during SCN (M=8.58, SD=4.42), followed by DF (M=8.5, SD=3.7) and lowest for OB (M=7.08, SD= 4.38). Post hoc tests revealed that SCN elicited more fixation counts than OB (p = 0.016). Saccade counts followed a similar pattern with OB (M=173.9, SD= 57.4) eliciting fewer saccades than SCN (M=186.15, SD= 57.3, p = 0.04) and DF (M=200.95, SD= 60.45, p = 0.02). The opposite effects were found for fixation durations. That is, saccade durations were significantly longer during OB (M = 1762.8 ms, SD = 2085.7) compared to SCN (M = 1323.3 ms, SD = 872.6, *p* =.005) and DF (M = 1115.4 ms, SD = 693.0, *p* =.001). This pattern fits well in line with other studies showing that the visual exploration of scene-imagery elicits more but shorter fixations than object construction (McCormick & Maguire, 2021; Taube et al., 2025, preprint).

### 3.2 fMRI behavioural results

Participants performed successfully across all three conditions, F(2, 18) = 1.55, p =.239. Mean success ratings were M=1.35 (SD = 0.26) for SCN, M=1.31 (SD = 0.25) for OB, and M=1.52 (SD = 0.53) for DF.

### 3.3 Univariate fMRI results

For the contrast SCN > DF, we found strong activation in our hypothesized areas, namely bilateral vmPFC, bilateral medial temporal lobes, such as the parahippocampal gyri and extending into the hippocampi, as well as bilateral lateral and medial parietal cortices (see Fig. 2, and Table S1 for MNI coordinates). The opposite effect revealed predominantly activation in the language related left temporal and frontal regions, as well as in bilateral occipital cortices.

**Figure 2.**
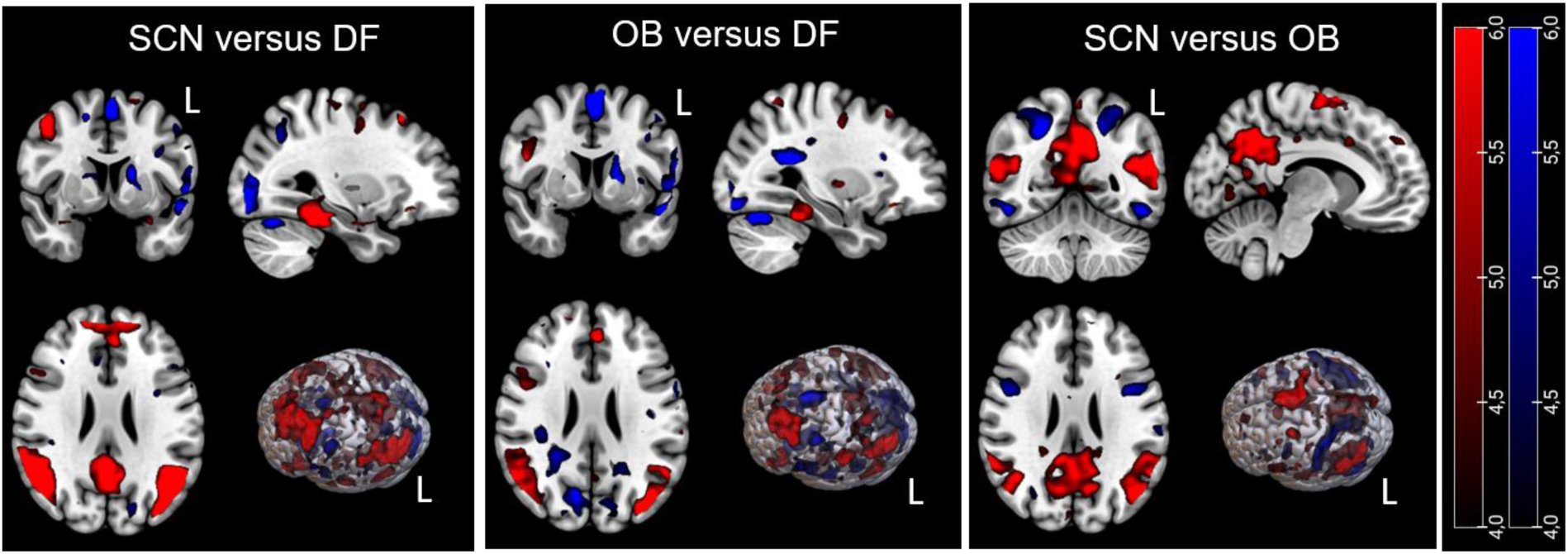
Group analysis of 7 Tesla fMRI standard whole-brain activation for scenarios, objects and definitions. Activations are thresholded at p<0.001 uncorrected and a cluster threshold at 10 adjecent voxels. T-maps are displayed on a T1-weighted MRI (MNI template), L=left.

In addition, when contrasting OB > DF, we found a very similar pattern, involving bilateral vmPFC, and parahippocampal gyri, as well as lateral and medial parietal cortices. For OB > DF, the hippocampus was not part of the pattern. Again, the opposite contrast, DF > OB, revealed a left-lateralized language pattern, including the lateral temporal and frontal cortices, as well as activation in bilateral occipital cortices.

Lastly, SCN resulted in higher activation of the lateral and medial parietal cortices than OB. Activation for SCN > OB in the vmPFC and medial temporal lobes, particularily parahippocampal gyri and right anterior hippocampus were only visible when lowering the threshold to a more liberal threshold of p=0.005. For the opposite contrast, we found higher bilateral activation in the inferior parietal cortex, and inferior temporal cortex for OB > SCN. No sub-threshold activation was found in the medial temporal lobes.

### 3.4 Multivoxel pattern analysis results

Next, we used anatomical ROIs to examine classifier accuracy in specific brain areas (see Fig. 3). For the vmPFC, accuracy exceeded chance only for SCN (M = 61.391, SD = 39.4, *t*(17) = 3.1, *p* = 0.01) but not for DF (M = 42.22, SD = 31.35, *t*(18) = 1.3, *p* = 0.23) or OB (M = 40.28, SD = 30.51, *t*(18) = 1.01, *p* = 0.33). In addition, we found that the classifier accuracy in the vmPFC exceeded the accuracy (F(2, 18) = 3.71, p=0.04). In posthoc analysis, SCN were classified with greater accuracy than OB (p = 0.02), and at a trend level DF (p = 0.05).

**Figure 3.**
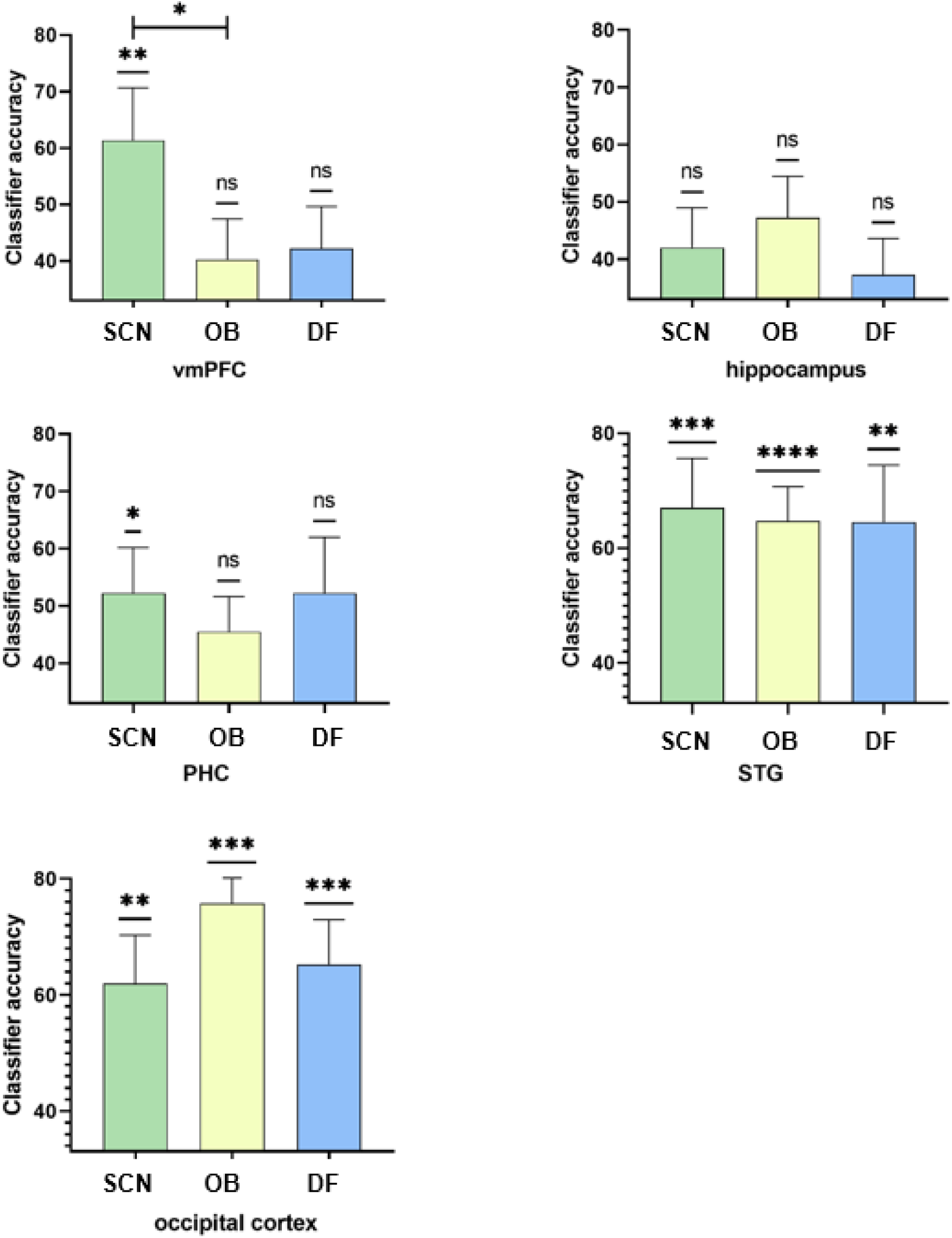
Classifier accuracies for the individual ROIs. For most ROIs, classifier accuracy exceeded chance level (33%, one-sample T-Tests, significance indicated above individual bars). Only the vmPFC showed differences between the conditions (confirmed by rm-ANOVA). Here, SCN was classified more often correctly than OB, and DF at a trend level. ^#^ = p<0.1 * = p<0.05, ** = p< 0.01, *** = p< 0.001, **** = p< 0.0001

For the parahippocampal gyrus, also only SCN accuracy exceeded chance (M = 52.25, SD = 35.56, *t*(19) = 2.421, *p* = 0.0257) but not for DF (M = 52.25, SD = 43.54, *t*(19) = 1.977, *p* = 0.0627) or OB (M = 45.50, SD = 27.43, *t*(19) = 2.038, *p* = 0.0557). However, there were no differences between conditions (F(2,18) = 0.45, p = 0.61).

For the hippocampus, accuracy did not significantly exceed chance for any condition: SCN (M = 42.00, SD = 31.05, *t*(19) = 1.296, *p* = 0.21), OB (M = 47.25, SD = 32.42, *t*(19) = 1.966, *p* = 0.064), and DF (M = 37.25, SD = 28.35, *t*(19) = 0.67, *p* = 0.51). Further, there were no differences between conditions (F(2,18) = 1.01, p = 0.36).

For the superior temporal gyrus (STG), accuracy significantly exceeded chance for all conditions: DF (M = 64.50, SD = 44.54, *t*(19) = 3.163, *p* = 0.0051), OB (M = 64.75, SD = 26.53, *t*(19) = 5.352, *p* < 0.0001), and SCN (M = 67.00, SD = 38.54, *t*(19) = 3.945, *p* = 0.0009), but no differences were found between the conditions (F(2,18) = 0.03, p = 0.96).

Similarly, accuracy within the occipital cortex significantly exceeded chance for all conditions: DF (M = 65.25, SD = 34.54, *t*(19) = 4.175, *p* = 0.0005), OB (M = 75.75, SD = 46.15, *t*(19) = 4.143, *p* = 0.0006), and SCN (M = 62.00, SD = 37.22, *t*(19) = 3.485, *p* = 0.0025), but no differences between conditions were found (F(2,18) = 0.81, p = 0.45).

As a control, we conducted an MVPA analysis on the whole brain and found that classifier accuracy significantly exceeded chance for all conditions with great accuracies: SCN (M = 83.00, SD = 43.66, *t*(19) = 5.121, *p* < 0.0001), OB (M = 73.50, SD = 40.30, *t*(19) = 4.495, *p* =0.0002), and DF (M = 73.25, SD = 47.52, *t*(19) = 3.788, *p* =0.0012). However, on the whole brain level, no differences in between conditions were found (F(2,18) = 0.48, p = 0.62), indicating no general neural differences between conditions.

### 3.5 After scanning task results

Upon visual inspection, all participants completed the drawings in an effortful manner, indicating a good task compliance (see Fig. 4 for an example). The debriefing ratings were on average very high for all conditions (SCN: 4.4 ± 1.01, OB 4.5 ± 0.6, and DF 4.5 ± 0.8). A two-way repeated measures ANOVA revealed a significant main effect of “Rater” (F(1, 19) = 6.31, *p* =.02), indicating that Rater 1 assigned higher values than Rater 2, but no significant interaction was found between “Rater” × “Condition” (*F*(2, 38) = 2.95, *p* =.06). Also, the reliability between the two raters showed a strong correlation for all conditions (Spearman’s correlation, SCN ρ=0.705, p=0.010, OB ρ=0.529, p=0.050, and DF ρ=0.919, p=0,01).

**Figure 4.**
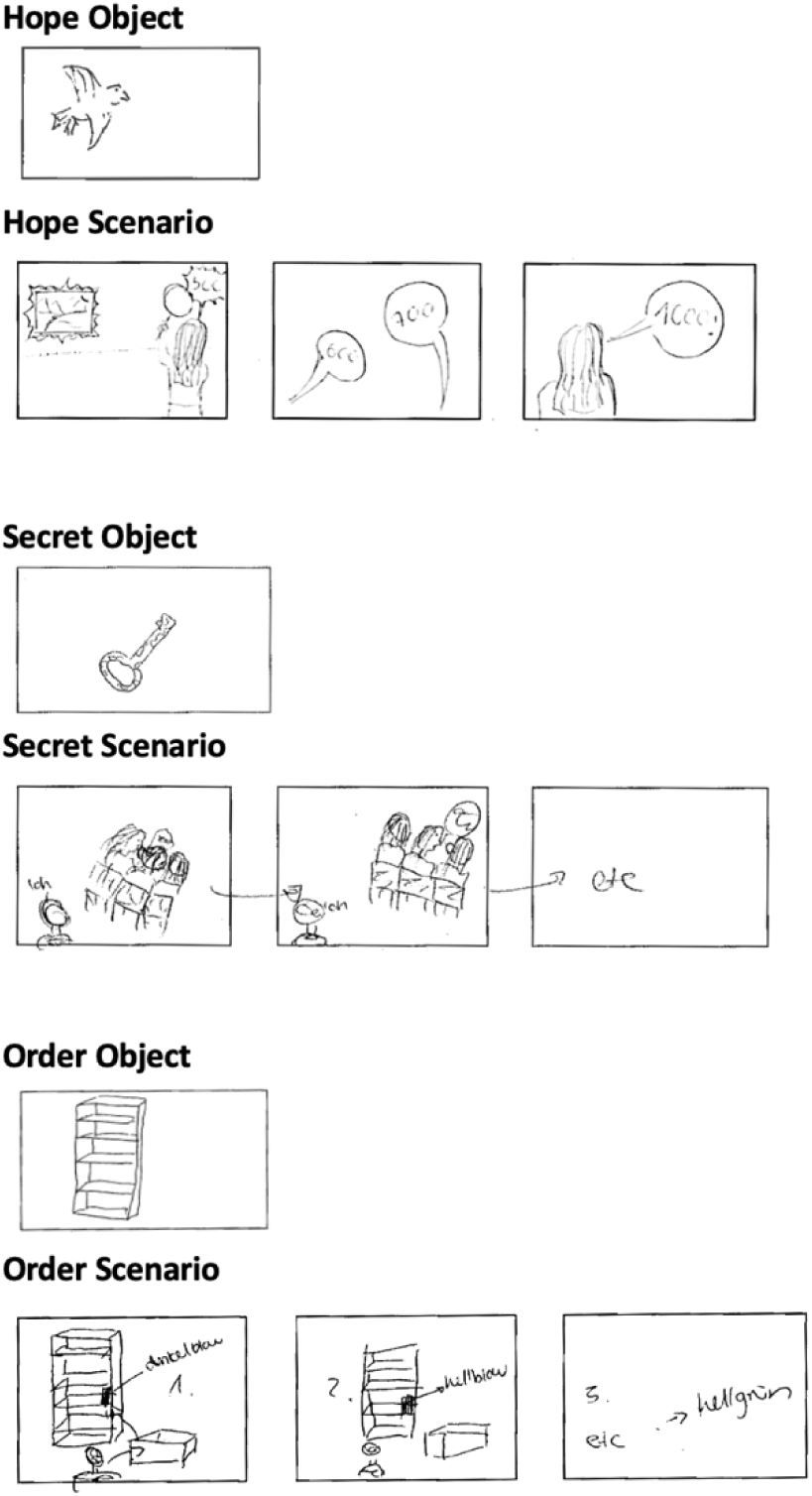
Representative drawings from a participant. Drawings were evaluated for clarity of representation, richness of detail, internal consistency, and perceived effort.

## Discussion

This study investigated the neural representation of mental imagery under varying degrees of perceptual richness and conceptual abstraction. Specifically, we asked whether the ventromedial prefrontal cortex (vmPFC), traditionally implicated in schematic and conceptual processes, also represents perceptually vivid imagery content. By directly comparing three tasks, scenario construction (SCN), object construction (OB), and abstract definitions (DF), we sought to disentangle schematic abstraction from visual richness. Our findings suggest a specific representational role for the vmPFC, advancing theoretical models of scene and scenario construction and shedding light on the neural underpinnings of imagery variability, including phenomena like aphantasia.

In line with previous literature (Ciaramelli et al., 2019; Gilboa & Marlatte, 2017; McCormick, Ciaramelli, et al., 2018), our univariate results demonstrate that the vmPFC was more strongly activated during SCN and OB conditions compared to DF. This reinforces the notion that vmPFC engagement is not restricted to abstract conceptual processing but also, and maybe exceedingly, occurs during perceptually rich mental simulation. Activation of the vmPFC during both SCN and OB (vs. DF) conditions supports theories suggesting its involvement in scenario construction and imagination (McCormick & Maguire, 2021) possibly by integrating schematic structures with perceptual content (Monk et al., 2021). Importantly, stronger vmPFC activation for SCN compared to OB emerged at more liberal thresholds, suggesting a graded sensitivity to content richness.

Critically, MVPA revealed that vmPFC classifier accuracy was significantly above chance only during scenario construction with classification accuracy for SCN exceeding OB and at a trend level of p=0.05 for DF within this region. This finding demonstrates that the vmPFC encodes distinct, distributed neural patterns specifically associated with imagery-rich scenarios, extending beyond general schema-guided processes. These results align with previous evidence showing that specific video clips could be identified within the vmPFC using pattern classification, with accuracy correlating positively with subjective vividness ratings (St-Laurent et al., 2015). Together, this evidence supports the dual-function model for the vmPFC, wherein this region not only orchestrates schema-guided retrieval (Ghosh & Gilboa, 2014; Monk et al., (Delg020) but also maintains content-specific representations, particularly when rich visual-spatial information is involved. This evidence is especially compelling, since MVPA captures representational fidelity beyond mean activation (Norman et al., 2006); thus directly supporting our hypothesis that the vmPFC contains fine-grained information about perceptually detailed mental content.

Notably, in medial temporal areas, our univariate data revealed robust activation of the hippocampus (HC) and parahippocampal cortex (PHC) during SCN > DF, and PHC (but not HC) during OB > DF. This aligns with previous research implicating the HC in binding elements into coherent scenes (Dalton et al., 2018; Hassabis & Maguire, 2009; Maguire & Mullally, 2013) and the PHC in representing spatial and contextual information (Aminoff et al., 2013; Epstein, 2008). The lack of significant hippocampal activation for OB > DF and the subthreshold HC effects for SCN > OB are consistent with the idea that full-fledged scene construction, as required for SCN, is necessary to robustly engage the HC (Clark et al., 2018; Dalton et al., 2018).

In contrast, MVPA in medial temporal lobe regions yielded more ambiguous results. In the PHC, classifier accuracy was above chance only for SCN but did not differ significantly between conditions. In the HC, accuracy did not exceed chance for any condition. These findings may reflect limitations in detecting distributed representations in these regions due to signal-to-noise constraints at high field strength or task demands that elicited less consistent spatial coding than canonical scene construction tasks (e.g., single scene snapshots rather than dynamic spatial integration).

The superior temporal gyrus (STG), often implicated as a semantic hub (Chadwick et al., 2016; Pobric et al., 2007), showed above-chance classifier accuracy for all three conditions, without significant differences. Since all conditions rely on some degree of semantic knowledge, this finding may reflect the idea that the STG is involved in generalized semantic elaboration, irrespective of imagery vividness (McCormick & Maguire, 2021; Monk et al., 2020). Similarly, occipital cortex classification was above chance across conditions, reflecting shared recruitment of visual-perceptual regions even during abstract or conceptual tasks (Cui et al., 2007; Kim et al., 2009; Naselaris et al., 2015).

Eye-tracking data further supported our hypothesis that imagery conditions differ in visual attention dynamics. As expected, participants exhibited more exploratory gaze behavior, indexed by saccade and fixation counts, during scenario construction, and longer fixations during object imagery. These findings which have also previously been reported (El Haj & Lenoble, 2018; McCormick & Maguire, 2021; Taube et al., 2025, preprint) align with the idea that scene-based imagery engages dynamic spatial exploration, while object imagery involves detailed scrutiny of a single element.

While we carefully designed our task to contrast scenario, object, and definition-based imagery using identical abstract nouns as cues, we acknowledge that isolating these mental processes with absolute precision is inherently challenging. Mental imagery is multifaceted, and participants may have engaged in overlapping strategies across conditions—for example, bringing in some object details while imagining definitions, or evoking schematic structures even during object imagery. However, we believe the converging evidence from multiple modalities, behavioral ratings, eye-tracking, univariate activation, MVPA, and post-scan debriefing, provides strong construct validity for our task differentiation.

Taken together, our results provide converging evidence that vmPFC contributes to the representation, not just the coordination, of perceptually rich, temporally extended mental scenarios. This challenges narrow views of vmPFC as solely a schematic or abstract hub (devoid of details) and supports integrative models that position it as a central node in a distributed imagery network. The unique role of the vmPFC in MVPA, compared to other regions such as the HC or PHC, highlights its sensitivity to the qualitative richness of imagined scenarios.

In the context of aphantasia, these findings underscore the neural distinctiveness of imagery-rich mental content and suggest that individuals lacking visual imagery may exhibit reduced representational specificity in vmPFC during simulation tasks, a hypothesis that could be tested in future work. More broadly, our study offers a high-resolution glimpse into how the brain represents internal experiences, bridging the gap between conceptual structure and perceptual content in imagination.

## Acknowledgements and funding

We thank all participants for their time and effort. This research was supported by the Hertie Network of Excellence in Clinical Neuroscience. Work in C.M.’s lab is further financed by internal research funding of the Faculty of Medicine (BONFOR), University Hospital Bonn, and by the Deutsche Forschungsgemeinschaft (DFG, German Research Foundation, MC 244/3-1). This project is further funded by the Federal Ministry of Research, Technology and Space (BMFTR) under the funding code (FKZ): 01EO2107.

---

**Table S1.** Whole brain group analysis.

| Region | x | y | z | T | Z | Cluster size (voxels) |
| --- | --- | --- | --- | --- | --- | --- |
| **EV > DF** |  |  |  |  |  |  |
| Precuneus | -6 | -54 | 40 | 13.73 | 6.67 | 46219 |
| supramarginal gyrus | -49 | -53 | 24 | 10.87 | 6.06 | 23030 |
| superior temporal gyrus | 56 | -54 | 16 | 10.13 | 5.87 | 22244 |
| *HC | 30 | -30 | -10 |  |  |  |
| middle frontal gyrus | -39 | 23 | 42 | 11.11 | 6.12 | 32292 |
|  | 55 | 12 | 36 | 8.27 | 5.32 | 4674 |
|  | 52 | 35 | 18 | 7.42 | 5.03 | 5008 |
| **DF > EV** |  |  |  |  |  |  |
| lingual gyrus | 20 | -92 | -2 | 8.30 | 5.33 | 8181 |
| cuneus | -10 | -96 | -3 | 7.57 | 5.08 | 5822 |
| **OB > DF** |  |  |  |  |  |  |
| middle frontal gyrus | -41 | 24 | 44 | 10.28 | 5.91 | 10744 |
| angular gyrus | -42 | -78 | 31 | 9.48 | 5.70 | 9554 |
| superior temporal gyrus | 54 | -59 | 27 | 5.55 | 4.23 | 10123 |
| postcentral gyrus | -34 | -41 | 53 | 5.31 | 4.11 | 5056 |
| inferior frontal gyrus | 48 | 15 | 37 | 5.26 | 4.08 | 4037 |
| **DF > OB** |  |  |  |  |  |  |
| superior frontal gyrus | -2 | 6 | 36 | 9.98 | 5.38 | 5695 |
| cuneus | 16 | -96 | 5 | 8.58 | 5.43 | 30297 |
| superior temporal gyrus | -54 | 9 | -12 | 7.06 | 4.89 | 3658 |

| EV > OB |  |  |  |  |  |  |
| --- | --- | --- | --- | --- | --- | --- |
| Precuneus | -7 | -55 | 42 | 10.91 | 6.07 | 35486 |
| medial Temporal lobe | -52 | -68 | 23 | 8.37 | 5.36 | 16743 |
| temporal lobe | 48 | -37 | -4 | 8.14 | 5.28 | 15476 |
| *HC | 30 | -30 | -10 |  |  |  |
| superior frontal gyrus | -5 | 8 | 69 | 7.37 | 5.00 | 8616 |
| OB > EV |  |  |  |  |  |  |
| middle occipital gyrus | -51 | -68 | -7 | 8.08 | 5.26 | 4214 |
| inferior parietal lobe | -43 | -37 | 48 | 7.18 | 4.94 | 7073 |
| precuneus | 24 | -54 | 46 | 6.93 | 4.84 | 6930 |

P < 0.001 uncorrected, Voxel cluster 10 adjacent voxels
\*part of a greater cluster

## Notes

### Competing Interest Statement

The authors have declared no competing interest.

